# Intracellular barriers and receptor masking limit success of a *Pseudomonas aeruginosa* clinical phage family

**DOI:** 10.64898/2026.06.18.732992

**Authors:** Wearn-Xin Yee, Amy B. Banta, Ryan D. Ward, Sriharshita Musunuri, Miaoxi Liu, Erin Huiting, Julia Gordeeva, Suzanne C. Letham, Tanmay A. M. Bharat, Jason M. Peters, Joseph Bondy-Denomy

**Affiliations:** Department of Microbiology & Immunology, University of California, San Francisco, CA, USA; Center for Genomic Science Innovation, University of Wisconsin-Madison, Madison, WI, USA; Structural Studies Division, MRC Laboratory of Molecular Biology, Cambridge, United Kingdom; Department of Medical Genetics, University of Wisconsin-Madison, Madison, WI, USA; Department of Bacteriology, University of Wisconsin-Madison, Madison, WI, USA; Department of Medical Microbiology and Immunology, University of Wisconsin-Madison, Madison, WI, USA; Great Lakes Bioenergy Research Center, University of Wisconsin-Madison, Madison, WI, USA

## Abstract

Bacteriophage therapy is needed to treat antibiotic resistant infections; however, when a clinical isolate resists a given phage, it is often unclear why. It is therefore currently unknown how to rationally fortify phage therapies to circumvent *a priori* resistance. Using a family of broad host range therapeutic *Pseudomonas aeruginosa* phages (*Pbunaviruses)*, we show that cell surface receptor masking and intracellular defenses are both common barriers in distinct clinical isolates. In some cases these barriers can be bypassed by intrafamily phage engineering. Using unbiased genome-wide CRISPRi screens, we reveal that the broadly conserved L-Rhamnose in the core polysaccharide is the receptor for *Pbunavirus* family. This molecule is often masked by diverse O-antigen structures. In other isolates with the L-Rha receptor accessible, internal defense mechanisms commonly prevent *Pbunavirus* DNA replication. A single anti-defense locus often encoding 8-11 different genes within the *Pbunavirus* family is required for optimal host range, providing anti-defense genes that enable replication of both *Pbunavirus* phages and phages of other families. Our work demonstrates the importance of both internal and surface defense mechanisms in clinical isolates causally antagonizing a commonly used phage therapeutic and presents phage engineering strategies to circumvent *a priori* resistance.

## Introduction

The widening gap between the growing burden of resistant infections and the shrinking arsenal of effective drugs^1,2^ has driven renewed interest in bacteriophage (phage) therapy as an alternative or adjunct to conventional antibiotics. Recent compassionate-use cases have demonstrated the clinical potential of phage therapy against difficult-to-treat infections caused by *Pseudomonas aeruginosa* and *Mycobacterium* species ^3,4^, among others. However, a central limitation of phage therapy relative to broad-spectrum antibiotics is the narrow host range of individual phages, which restricts their off-the-shelf utility and necessitates patient-specific phage selection or cocktail design.

*Pbunavirus* phages are among the most clinically deployed phages against *P. aeruginosa* due to their broad host range^5,6^. Members such as E217 and 14-1 are frequently included in cocktails with other phage families targeting distinct surface receptors to further broaden coverage^4,7,8^. Despite the therapeutic importance of this family, and a general acceptance of it being “LPS-specific”, the surface receptor recognized by Pbunaviruses has not been resolved. Recent studies have demonstrated that Pbunaviruses also encode inhibitors of phage defense systems which, when expressed *in trans*, sensitize bacteria to both related and unrelated phages^9,10^. Together, these findings suggest that the *Pbunavirus* family harbors a rich repertoire of evolutionary strategies for entering and killing diverse *P. aeruginosa* strains. The central question motivating this work is therefore: when a *Pbunavirus* phage fails against a clinical isolate, what barriers are responsible, and can they be overcome through intrafamily engineering?

The discovery of numerous antiphage defense systems in bacterial genomes has raised the possibility that these systems limit phage host range, particularly given their frequent localization within variable accessory genomes^11,12^. However, most defense systems have been characterized through heterologous expression, and their functional contribution to phage resistance in clinical isolates remains poorly understood^13^. Machine-learning approaches have begun to bridge this gap by generating large-scale phage-host interaction matrices, but these studies predominantly implicate surface barriers rather than intracellular defense systems as the primary determinants of phage susceptibility^6,14,15^. Systematic approaches are needed that combine experimentally validated phage-host susceptibility data with targeted phage engineering to identify and overcome specific barriers to infection. This approach can also identify any potential trade-offs to phage modifications that may limit host range.

Here, we dissect the barriers that prevent *Pbunavirus* phages from infecting clinical isolates and ask whether these barriers can be systematically identified and overcome. Testing eight Pbunaviruses against 39 diverse clinical strains, we find that *a priori* resistance is explained by many distinct mechanisms, only some of which are naturally solved by the *Pbunaviruses*. Multiple distinct surface and intracellular barriers each operate in a substantial and largely non-overlapping fraction of isolates. We identify the conserved *Pbunavirus* receptor (L-Rhamnose), characterize the environmental conditions that govern its accessibility, and define a phage-encoded anti-defense locus whose deletion unmasks intracellular defenses that are otherwise neutralized. However, some intracellular barriers that defeat wild-type phages remain unresolved, defining a clear limitation of the current *Pbunavirus* arsenal. We propose that solutions to strains with active intracellular barriers would require either additional sampling of more *Pbunavirus* phages, or sampling of phages from outside the *Pbunavirus* family to identify active anti-defense genes. For the strains where phages can breach both barriers and replicate, bacteria evolve phage resistance that selects for the loss of the core polysaccharide, which produces a stereotyped and therapeutically exploitable outcome that carries direct implications for how *Pbunavirus* phage treatment should be sequenced within multi-phage regimens with antibiotics.

## Results

### Identification of the *Pbunavirus* receptor

To dissect the mechanisms of *a priori Pbunavirus* resistance in *P. aeruginosa,* we began with the model laboratory strain PA14, originally isolated from a clinical source, which is generally resistant to *Pbunavirus* phages. PA14 harbors numerous known phage defense systems, including Jumbo phage killer, Gabija, Type I-F CRISPR-Cas, and Shango^16^. Given the density of defense systems in PA14, we used an unbiased approach to determine the causal barrier(s) that restrict *Pbunavirus* infection in this strain. We constructed a CRISPRi library targeting all genes in PA14 with multiple guide RNAs per gene, then challenged the library with phages F8 and 14-1, neither of which forms plaques on wild-type PA14. While F8 infection remained unsuccessful even at high multiplicities of infection (MOI of 1000), partial infection was observed with 14-1, enabling identification of strain dropouts via sequencing of guide RNAs. This analysis identified genes whose knockdown rendered bacteria susceptible to phage killing, revealed two loci: *waaL* and PA4011 (Figure 1A). CRISPRi guides targeting PA4011 only minimally enhanced 14-1 plaquing, but knockdown of *waaL* conferred robust sensitivity of PA14 to both F8 and 14-1. This result was independently corroborated using the PA14 ordered transposon library, where transposon insertion early within the *waaL* coding sequence similarly rendered PA14 susceptible to both 14-1 and F8 (Figure 1B). Due to the modest effect of knocking down PA4011, this gene was not followed up.

**Figure 1:**
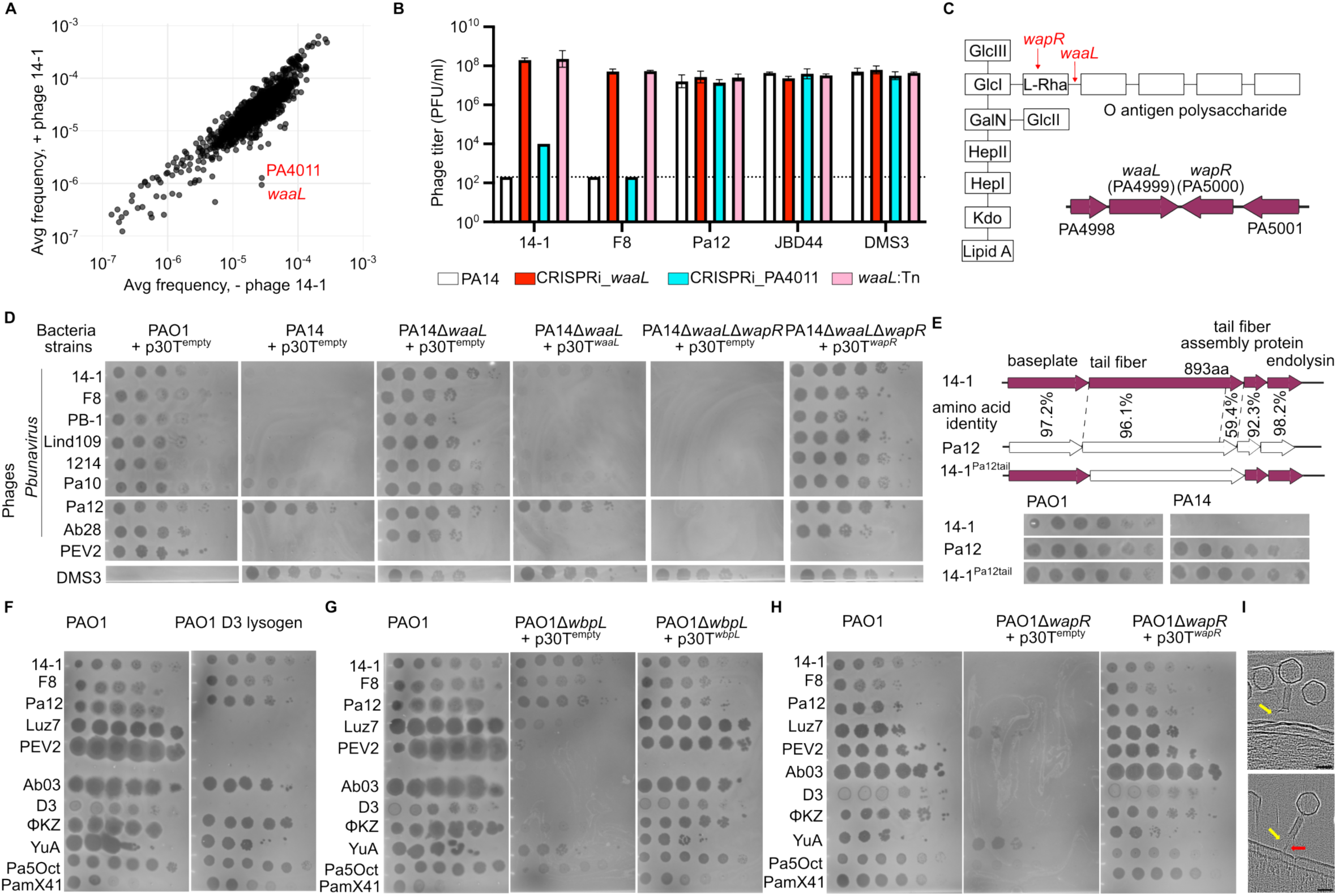
L-Rhamnose in the core LPS is the receptor for *Pbunavirus* phages. (A) Frequency of genes knocked down using CRISPRi with/without the addition of 14-1 phage. (B) Phage titer across different phages in the presence/absence of functional *waaL*. (C) Graphical representation of the role of *waaL* and *wapR* in O-antigen lipopolysaccharide synthesis; genetic locus of waaL and wapR is as shown. (D) Plaque assay of different phages in PA14 with Δ*waaL* with/without Δ*wapR*, as well as the respective complementation. (E) Schematic of tail fiber swap between 14-1 and Pa12 and plaque assays on PA14. Amino acid identity is calculated using ClustalOmega. (F) Plaque assay of different phages on PAO1 and D3 lysogen. (G) Plaque assay of different phages on PAO1, PAO1Δ*wbpL* with/without complementation. (H) Plaque assay of different phages on PAO1, PAO1Δ*wapR* with/without complementation. (I) Two slices from cryo ET data of 14-1 phage infecting pAO1 cells. Yellow: phage tail fibers extending close to the outer membrane. Red: phage sheath orthogonal to the outer membrane, presumably in the act of injection. Scale bar: 50nm.

The *P. aeruginosa* cell surface displays three polysaccharide structures: the common polysaccharide antigen (CPA), the O-specific antigen (OSA) and the uncapped LPS. PA14 lacks CPA and the uncapped LPS owing to a defective glycosyltransferase *wbpX* and *migA* gene respectively, and so possesses only OSA^17^. WaaL is an O-antigen ligase that attaches the O-antigen to the L-Rha residue of the LPS core^18^ (Figure 1C). Deletion of *waaL* removes the O-antigen, exposing only the core LPS, including the L-Rha residue, on the PA14 surface. Consistent with above, PA14Δ*waaL* becomes susceptible to all other *Pbunavirus* phages tested. However, further deletion of *wapR*, which encodes the rhamnosyltransferase responsible for adding L-Rha to the core, abolished susceptibility to all *Pbunavirus* phages. Complementation *in trans* restored the expected phenotypes (Figure 1D). Pa12 showed reduced adsorption to PA14Δ*waaL*Δ*wapR* which is similarly complemented *in trans* (Figure S1A). These results establish L-Rha on the core LPS as the receptor for the *Pbunavirus* family and demonstrate that the receptor is present but masked by the O-antigen in PA14, a resistant strain.

### A phage tail fiber swap overcomes receptor masking

The observation that Pa12 forms plaques on PA14, despite the L-Rha receptor masking, suggested that Pa12 possesses a tail protein capable of penetrating the O-antigen barrier to reach the underlying core. In a related phage E217, structural studies have shown that the N-terminal domain (NTD) of the tail fiber (gp45 in 14-1) mediates binding to the baseplate, while the C-terminal domain (CTD) extends outward to contact the bacterial surface^19^. The tail fiber has been directly implicated in variable receptor access among Pbunaviruses: evolution of *Pbunavirus* phages for enhanced biofilm targeting yielded amino acid substitutions specifically within the tail fiber^20^. We hypothesized that natural diversity within the *Pbunavirus* family could contain solutions to surface barriers. Comparing Pa12 and 14-1 tail fiber revealed high conservation in the NTD (96.1% amino acid identity) and high divergence in the C-terminal 69 amino acids (59.4% amino acid identity) (Figure 1E). We therefore constructed a chimeric phage carrying the Pa12 tail fiber on the 14-1 backbone (14-1^Pa12tail^) and tested it alongside parental 14-1 and Pa12. The chimeric phage formed plaques on PA14, whereas parental 14-1 did not, demonstrating that a single tail fiber swap can overcome receptor masking. This result establishes that natural tail fiber diversity within the *Pbunavirus* family provides ready-made solutions to the PA14 surface barrier, and that rational tail fiber engineering between related phages represents a practical strategy for broadening host range.

### L-Rha defines a distinct receptor class among LPS-dependent phages

Remarkably, many *P. aeruginosa* phages are described as LPS-dependent, but this is seldom defined more specifically. To determine if L-Rha is a common receptor among other LPS-dependent phages, we tested phage susceptibility across a series of PAO1 surface mutants. A D3 lysogen, which modifies the O-antigen structure^21^, inhibited D3, LUZ7, and PEV2 but did not decrease the efficiency of plating (EOP) of the *Pbunavirus* phages or other LPS-dependent phages such as Ab03 (*Nankokuvirus*). Pilus/flagellum-dependent phages also remained unaffected by D3 lysogeny (Figure 1F). Deletion of *wbpL*, which is required to initiate synthesis of the repeating units of both O-antigen and CPA in PAO1^22^ but not of the LPS core, abolished infection by all LPS-dependent phages except the Pbunaviruses, which remained fully capable of forming plaques. This is consistent with this family using L-Rha as opposed to any other element of the O-antigen structure. Complementation with *wbpL* expressed *in trans* restored susceptibility (Figure 1G). We also noted decreased plating efficiency for PaMx41 on Δ*wbpL*, suggesting that this phage likely uses LPS as its receptor. Together, these results separate phages that strictly require a specific O-antigen (D3, PEV2) from those that tolerate its modification or loss (Ab03, PaMx41, Pbunaviruses).

Having established that the Pbunaviruses are O-antigen-independent, we next asked whether they depend on the LPS core. Deletion of *wapR*, mirroring our PA14 results, abolished infection by both *Pbunavirus* and all other LPS-dependent phages (Figure 1H), consistent with our findings that Pbunaviruses recognize core LPS, specifically L-Rha, rather than any surface-distal elements extending from the cell envelope. This conclusion is further supported by cryo-ET imaging of *Pbunavirus* 14-1 phages infecting PAO1 cells (Figure 1I, S1B). Phages were observed at various stages of infection and cell entry. Some phages had their sheaths oriented orthogonal to the plane of the bacterial outer membrane, extending all the way to the outer membrane surface, presumably reflecting active injection of genetic material. Several other phages, however, were tilted away from an orthogonal orientation with respect to the outer membrane, with a single or several tail fibers extended toward and bound near the outer membrane, consistent with the location of the core oligosaccharides of the LPS^23^ (Figure 1I, S1B).

Interestingly, the flagellum- and pilus-dependent jumbo phage ϕKZ was also inhibited by loss of L-Rha in PAO1, unlike the pilus-dependent YuA. Loss of *wapR* inhibited not only ϕKZ but also other related jumbo phages (Figure S1C), suggesting that the *Phikzvirus* family uses core LPS as a secondary receptor. PAO1Δ*wapR*’s swimming or twitching motility remained intact, suggesting that the inability of ϕKZ to infect this strain is not due to loss of flagellum or pilus (Figure S1D). Collectively, these data reveal three distinct subgroups among “LPS-dependent” phages: those strictly dependent on a specific O-antigen (e.g. D3, PEV2), those tolerant of O-antigen modifications (e.g. Ab03, PaMx41), and those bypassing the O-antigen and requiring core LPS components, specifically L-Rha (e.g. the *Pbunavirus* and *Phikzvirus* families). Notably, phages in the latter two families have been used therapeutically and perhaps the use of a receptor that is ubiquitously present on any *P. aeruginosa* strain that possess an O-antigen^24^ endows broad host range.

### Nutrient-depleted media alter surface barriers

Apart from nutrient rich media (e.g. LB), the M9 minimal medium is often used experimentally to study biofilms and slow growth in *P. aeruginosa*^25,26^. We therefore tested *Pbunavirus* susceptibility in M9 minimal medium supplemented with 1% glucose. We observed that F8 efficiently lysed PAO1 in LB but not under minimal media conditions (Figure 2A).

**Figure 2:**
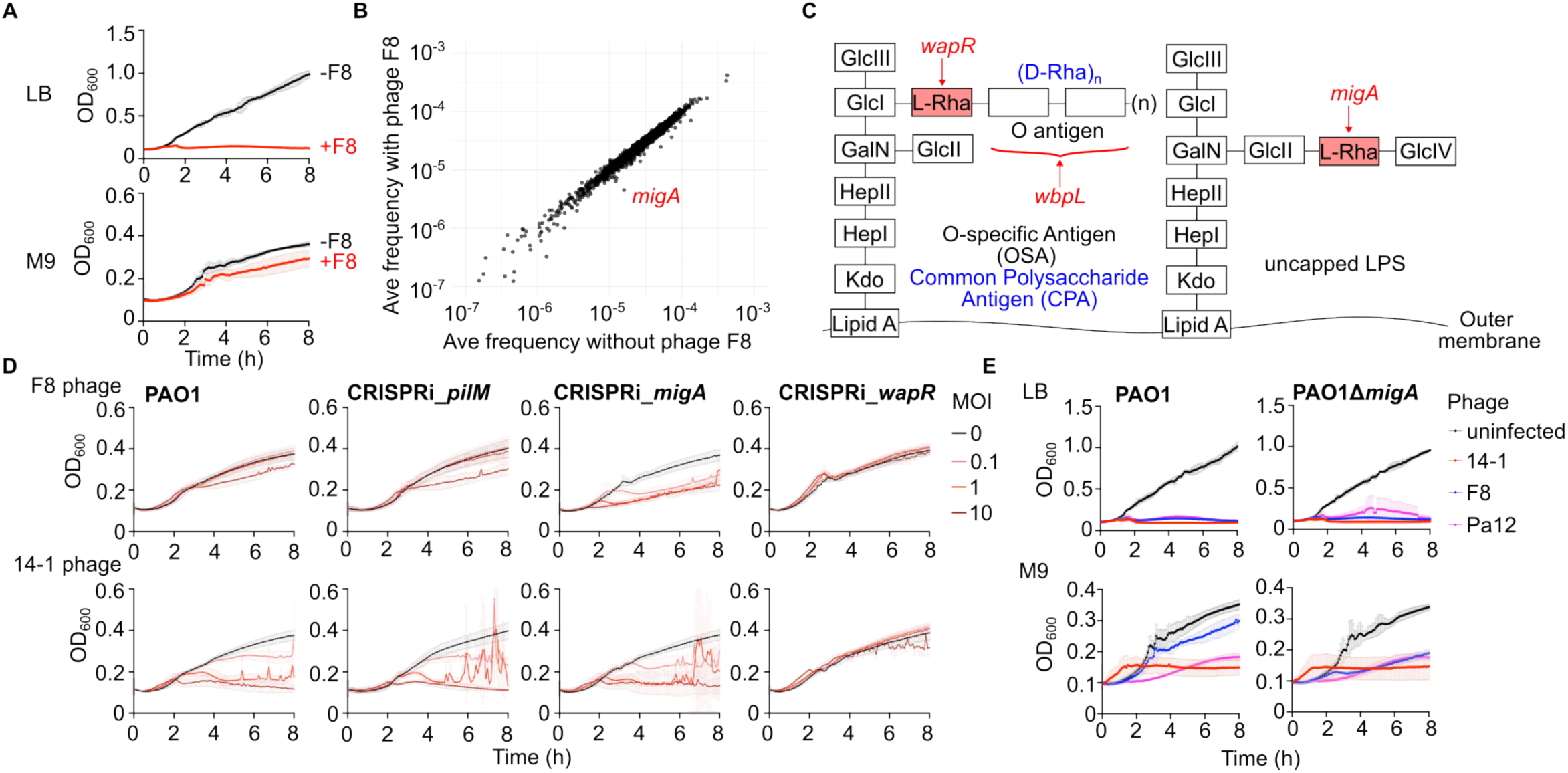
The uncapped LPS acts as a surface barrier against *Pbunavirus* phages in minimal media. (A) Growth curve of PAO1 under LB or M9 in the presence of 14-1 or F8. (B) Frequency of each gene knocked down using CRISPRi in the absence/presence of F8. (C) Graphical representation of the role of *wapR*, *wbpL* and *migA* in lipopolysaccharide (LPS) synthesis. (D) Growth curve of PAO1 with genes as indicated knocked down using CRISPRi under M9 conditions, in the presence of either 14-1 or F8 phages. (E) Growth curve of PAO1 and PAO1Δ*migA* with/without *Pbunavirus* phages in LB or M9.

To identify the causal factor inhibiting F8 infection in M9, we constructed and screened a whole-genome CRISPRi library in PAO1. Screening of the PAO1 library infected with F8 in M9 medium identified *migA* as the host factor conferring nutrient-limiting resistance (Figure 2B). MigA is a rhamnosyltransferase involved in uncapped LPS synthesis that competes with WapR for the L-Rha substrate and are differentially regulated with respect to each other^27,28^(Figure 2C). Due to a mutation in *migA*, the uncapped LPS is absent in PA14^17^ but present in PAO1. In minimal media, increased uncapped LPS production likely masks the L-Rha receptor, preventing F8 tail fibers from reaching their target. To confirm the role of *migA*, we inhibited *migA* using a CRISPRi guide in PAO1, with guides against *wapR* and *pilM* as positive and negative controls, respectively. 14-1, which lyses PAO1 in minimal media, was used as a control phage. Knockdown of *migA* increased susceptibility of PAO1 to F8 even at low MOIs (Figure 2D). A PAO1Δ*migA* deletion mutant similarly increased susceptibility to F8 specifically in M9 media, with no effect in LB or on phages 14-1 and Pa12, which already lyse PAO1 efficiently in M9 (Figure 2E). These results demonstrate that the uncapped LPS acts as a condition-dependent surface barrier in minimal media that can be overcome by *Pbunavirus* phages 14-1 and Pa12 and strongly caution that *in vitro* susceptibility in rich media may not predict therapeutic efficacy.

### Intracellular defense systems must be inhibited for success in clinical isolates

Having established the molecular identity of the conserved *Pbunavirus* receptor and the mechanisms by which surface structures can mask it, we next asked how these principles extend across distinct clinical strains. We challenged 39 *P. aeruginosa* isolates including laboratory strains PAO1, PA14, and PAK, multidrug-resistant MRSN isolates^29^, and clinical isolates from geographically diverse sources with eight *Pbunavirus* phages and two control phages from different families. By calculating efficiency of plating relative to PAO1, we found that 16 were susceptible to at least one *Pbunavirus*, and 7 were susceptible to all, confirming that Pbunaviruses possess a comparatively broad but incomplete host range (Figure 3A-B). Genome sequencing and analysis confirmed that these isolates carry a variety of known antiphage defense systems. The number of defense system carried by isolates that are susceptible to at least one *Pbunavirus* phage was not statistically different compared to the number of defense systems carried by isolates resistant to all *Pbunavirus* phages (*p* = 0.1613), suggesting that in our dataset, no direct correlation between number of defense systems and *Pbunavirus* susceptibility was observed.

**Figure 3:**
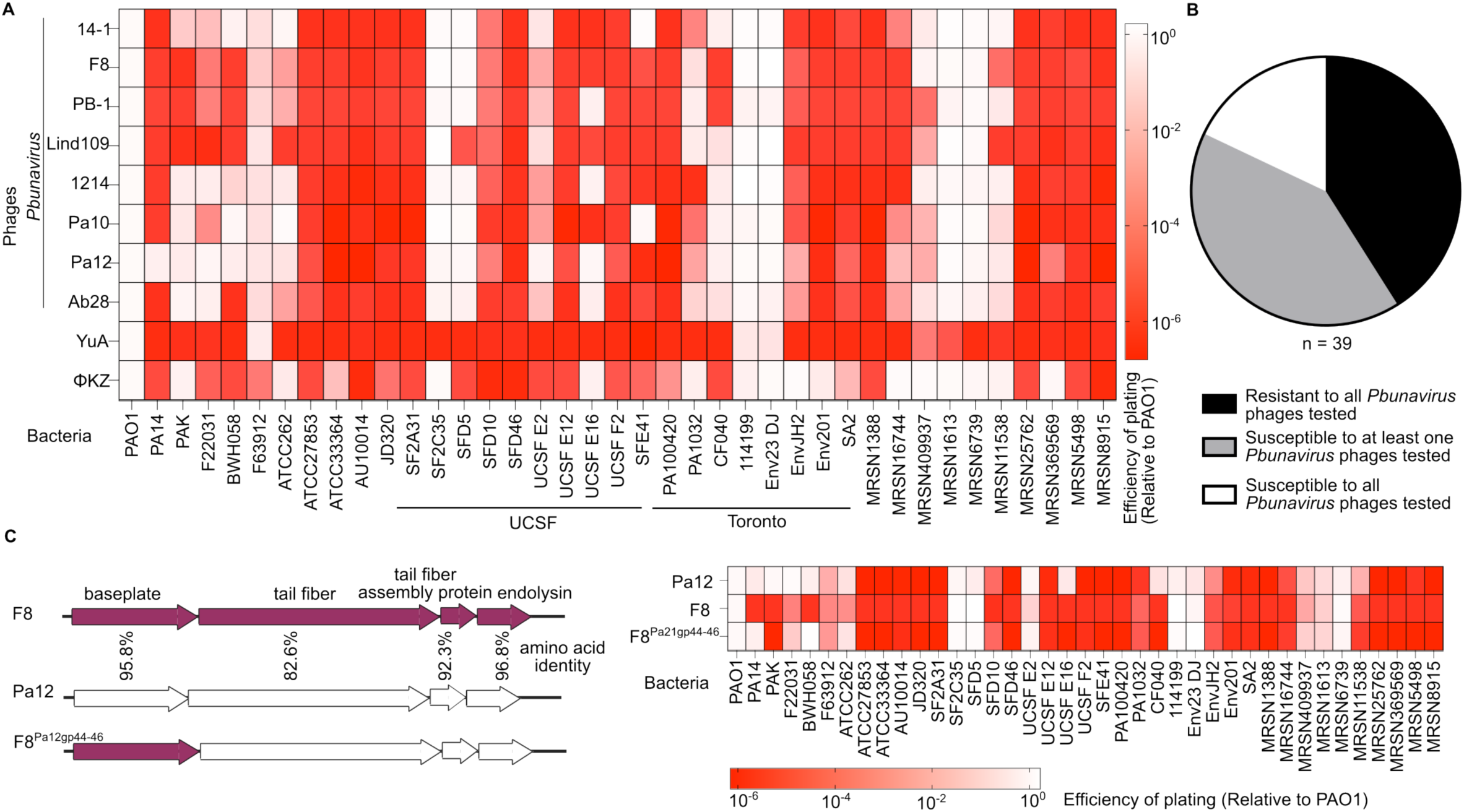
*P. aeruginosa* clinical isolates are differentially susceptible to *Pbunavirus* phages. (A) Efficiency of plating (EOP) of different strains relative to PAO1 of eight different *Pbunavirus* phages, as well as two phages from different families, YuA and ΦKZ. (B) Percentage of strains that are resistant to all *Pbunavirus* phages tested, susceptible to at least one *Pbunavirus* phages tested, or susceptible to all *Pbunavirus* phages tested. (C) EOP of different strains relative to PAO1 of F8, Pa12, and F8 with Pa12’s tail construct. Schematic of swap is as shown; amino acid identity is calculated using ClustalOmega.

These data raised the question of whether surface exclusion is the dominant mechanism by which *P. aeruginosa* excludes *Pbunavirus* infection. To address this, we first recombined Pa12’s tail fiber region into F8 and observed that this increased F8 host range by 4 strains, again suggesting that L-Rha masking is one mechanism to exclude members of this phage family (Figure 3C). However, we observed that this recombined phage was still unable to infect some clinical isolates that Pa12 and 14-1 can. Therefore, we asked which *Pbunavirus* phage gene(s) is required to succeed in these strains. Among the variably resistant isolates, inhibition of the END nuclease by gp88 from 14-1 (a gene encoded by only some *Pbunavirus* phages) enables success in strains CF040 and PAK^9^. Analysis of the F22031 genome showed the presence of the END nuclease. 14-1 lost the ability to plaque on F22031 upon gp88 inactivation (Figure 4A), confirming that END nuclease is also a causal *Pbunavirus* barrier in that strain.

**Figure 4:**
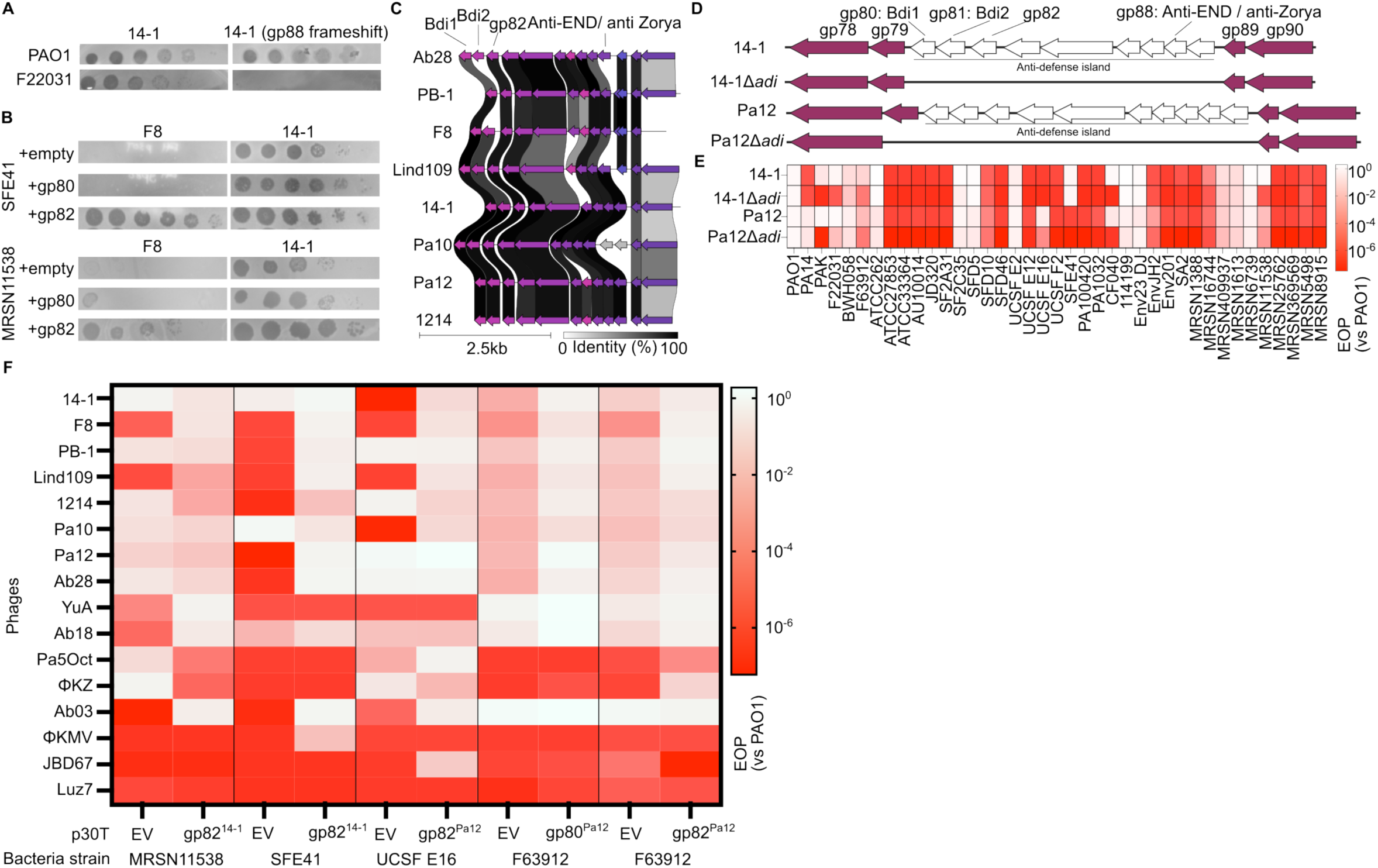
The pan-*Pbunavirus* genome encodes for anti-defenses which increases EOP of phages from multiple phage families on multiple clinical isolates. (A) Plaque assay of 14-1 wild type and with gp88 frameshift mutation on F22031 (B) Plaque assay of *Pbunavirus* phages on SFE41 or MRSN11538 with/without anti-defenses from 14-1 *in trans*. (C) Alignment of the anti-defense locus of *Pbunavirus* phages tested in this paper. (D) Schematic of deletion of anti-defense locus in 14-1 and Pa12. Anti-defenses Bdi1, Bdi2, anti-END/anti-Zorya previous identified^9,10^ are highlighted. (E) Efficiency of plating (EOP) of 14-1 and Pa12 with/without anti-defenses against different clinical isolates. (F) Heatmap showing EOP of phages from different families on F63912, UCSF E16, SFE41 and MRSN11538 in the absence/presence of anti-defenses identified in Pa12/14-1.

Among the remaining variably resistant isolates, MRSN11538 (a multidrug-resistant strain) and SFE41 were susceptible to 14-1 but not F8 (Figure 3A). To identify the 14-1 mechanism responsible, we screened previously constructed 14-1/F8 hybrid phages on these strains and mapped the activity to the gp80-82 region (Figure S3A). Individual cloning of gp80 and gp82 then revealed gp82 as a single gene that could rescue F8 phage replication on this strain (Figure 4B). Together with the independent identification of gp80 and gp81 as broad defense inhibitors (Bdi1 and Bdi2)^10^, these data define gp80-88 in 14-1 as an anti-defense island (“*adi*”). Comparative alignment of this region across our *Pbunavirus* collection revealed extensive gene gain, loss, and sequence divergence, highlighting the evolutionary plasticity of this locus (Figure 4C). This variation suggests that individual *Pbunavirus* phages have assembled distinct repertoires of anti-defenses against defense systems in clinical isolates.

To assess the functional importance of the anti-defense island, we deleted the entire *adi* region from both 14-1 and Pa12 (Figure 4D). Deletion did not increase host range, confirming that none of the putative anti-defenses inadvertently activate cryptic defense systems in our tested isolates. However, of the 18 strains initially susceptible to wild-type 14-1 and/or Pa12, 6 became resistant to the Δ*adi* mutants, including PAK, F22031, CF040 and MRSN11538 previously characterized (Figure 4E). We also observed that two strains F63912 and UCSF E16 became resistant to Pa12 upon deletion of the anti-defense island. Systematic cloning of anti-defense sub-regions again identified gp82^Pa12^ (a Pa12 variant of gp82^14–1^) as restoring Pa12Δ*adi* EOP on E16, and gp80^Pa12^ and gp82^Pa12^ as restoring activity against F63912 (Figure S3B). Expression of these anti-defenses *in trans* also induced complete susceptibility to all tested *Pbunavirus* phages, confirming that the internal defense targets of these anti-defenses are the primary barrier in these strains.

### *Pbunavirus* gp82 helps diverse phages replicate in clinical isolates

We next investigated whether anti-defense genes identified in 14-1 and Pa12 could increase EOP of *Pbunavirus* phages and phages from other families on clinical isolates. We tested nine additional phages spanning diverse families: YuA (*Yuavirus*), Ab18 (*Abidjanvirus*), Pa5Oct (*Wroclawvirus*), Ab03 (*Nankokuvirus*), JBD67 (*Beetrevirus*), ϕKMV (*Phikmvvirus*), LUZ7 (*Luzseptimavirus*), and ϕKZ (*Phikzvirus*). We observed increased replication of many distinct phages in MRSN11538 and SFE41 induced by gp82^14–1^, and similarly, gp82^Pa12^ improve replication of diverse phages in UCSF E16. These data suggest that gp82 and its homologues target a defense system or systems that are broadly active (Figure 4F, Figure S4). On F63912, gp82^Pa12^ not only increased EOP of the *Pbunavirus* phages, but also of ϕKZ. Despite gp82 neutralizing defense systems in all four strains, no singular defense system is shared between all four strains and yet not found on other strains (Figure S2), suggesting that gp82 is either a broad anti-defense, or that there is a defense system that has not yet been identified.

In contrast to gp82^Pa12^, gp80^Pa12^ did not enhance any other phage on F63912, a strain where it actively rescues *Pbunavirus* phages. These results demonstrate that *Pbunavirus* phages encode multiple anti-defenses within this locus (gp80/Bdi1, gp82, gp88/anti-END) and that gp82 and its homologues can broadly counter bacterial defenses, with therapeutic implications extending well beyond the *Pbunavirus* family itself.

### Intracellular defense systems in clinical isolates are mostly non-abortive

Of the 39 diverse *P. aeruginosa* isolates queried against the *Pbunavirus* collection, 16 were completely resistant to all phages tested (EOP <10^−4^). Although the mechanism of defense and a solution for this *a priori* resistance is not immediately clear, we asked whether phage failure was due to a surface-level or intracellular barrier. We performed single-step infection assays with Pa12, the broadest-range *Pbunavirus* in our collection of 39 isolates, with output phages titered on PAO1. Two strains (MRSN369569 and MRSN8915) were excluded because they produced phages that formed plaques on PAO1. Of 37 evaluable strains, Pa12 replicated productively in 16 (43%), failed to adsorb on 11 (30%), and adsorbed but failed to replicate in 10 (27%) (Figure 5A, Figure S5). These data suggest that intracellular defenses in clinical strains block *Pbunavirus* infection at a frequency comparable to surface barriers. Phage DNA replication was assessed in these 10 strains via qPCR, which revealed that 2 strains supported partial phage DNA replication while minimal DNA replication occurred in the remaining 8, suggesting that these defenses act at or prior to DNA replication (Figure 5B). This suggests that all defense systems in these clinical isolates likely inhibited Pa12 at or prior to phage DNA replication. Importantly, these data establish that intracellular defenses are not a rare exception but a pervasive and therapeutically relevant obstacle for *Pbunavirus* phages in clinical isolates.

**Figure 5:**
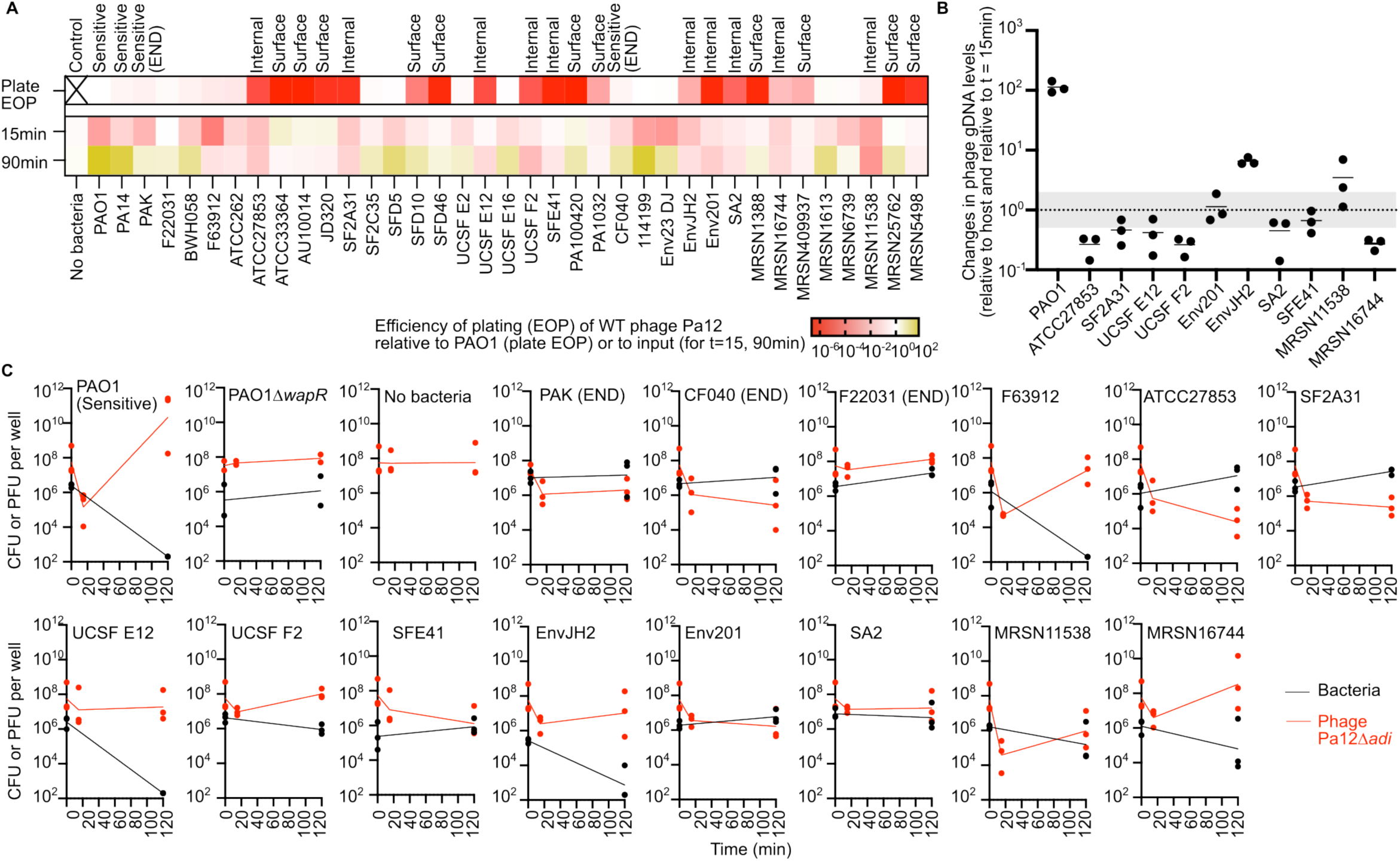
Pbunavirus phages face both internal and surface barriers; internal barriers inhibit phage DNA replication. (A) Single step growth curve of Pa12 phage infection across different clinical isolates, with PFU relative to input shown at t=15 and t=90min. Pa12’s plaque assay results from Figure 3A is included under “Plaque EOP” (B) qPCR to determine changes in phage gDNA levels at t=90 min compare to t=15 min. PAO1 was used as a positive control indicating phage replication. (C) Single step infection / growth curve of both Pa12Δ*adi* and different bacterial strains over time at MOIs 5 - 200.

To determine if active intracellular defenses lead to cell death, we used Pa12Δ*adi,* which we showed no longer carried anti-defenses compared to the parental Pa12 phage (Figure 4D, E). In addition to the 10 strains that we showed had an internal barrier mentioned above (Figure 5A, Figure S5), we included an additional 4 strains (PAK, F63912, F22031, CF040) that restricted Pa12Δ*adi* but not Pa12. Importantly, only 2/14 of these interactions excluding PAO1 (UCSF E12 and EnvJH2) led to cell death without phage replication, suggesting that most strains are antagonizing phage replication without enacting an abortive infection system (Figure 5C). This demonstrates that approximately one-third of susceptible clinical isolates carry therapeutically relevant internal defense systems that are actively neutralized by *Pbunavirus*-encoded anti-defenses, and that these anti-defenses are essential for productive infection. Additionally, most of the intracellular defenses did not display an abortive phenotype, i.e. the bacterium survives (Figure 5C). The future identification of the causal intracellular phage defenses and their circumvention is necessary to further broaden the host range of this phage family. These solutions may come from anti-defenses encoded by *Pbunavirus* isolates not tested here or derived from other phage families, the way *Pbunavirus* defense anti-defenses can rescue non-*Pbunavirus* phage families (Figure 4F, Figure S4). Alternatively, expression of bacterial toxins by *Pbunavirus* phages that gain access to the bacterial cytosol may enhance cell killing by these phages but would likely not endow phage replication.

### Evolved *Pbunavirus* resistance impairs cell fitness, slightly reduces antibiotic resistance but inhibits most phages

Having established an approach to identify and overcome some surface and intracellular barriers, with a long-term goal of overcoming all of them, we next asked whether successful *Pbunavirus* infection drives bacteria towards a consistent and exploitable resistance phenotype. We hypothesized that the L-Rha receptor in the core polysaccharide would need to be lost to achieve *Pbunavirus* resistance, which could compromise cell fitness, antibiotic sensitivity, and/or alter phage sensitivity. Previous studies have documented that *Pbunavirus* resistance frequently involves mutations in the LPS biosynthesis loci or large genomic deletions encompassing *galU*, a glucose-1-phosphate uridylyltransferase essential for core LPS biosynthesis^4,30–32^, the latter of which results in LPS truncation to the GalN residue (see Figure 1C for schematic)^33^. Additionally, different laboratories have reported variable changes in antibiotic sensitivity in response to *Pbunavirus* phage treatment^34,35^. Given that all Pbunaviruses rely on a single molecule (L-Rha) for infection, we asked whether its loss would alter the cell surface sufficiently to change cell fitness, antibiotic resistance or phage susceptibility in a consistent and predictable manner.

We evolved resistance in three genetic backgrounds: PAO1 (susceptible to both F8 and Pa12) and PA14 *waaL::Tn,* and PA14 with CRISPRi construct targeting *waaL* (Figure 6A), and determined the mutation leading to phage resistance, tested their growth, antibiotic susceptibility and susceptibility to other phage families. We observed that resistance can either be due to frameshift mutations in singular genes, or large chromosomal deletions.

**Figure 6:**
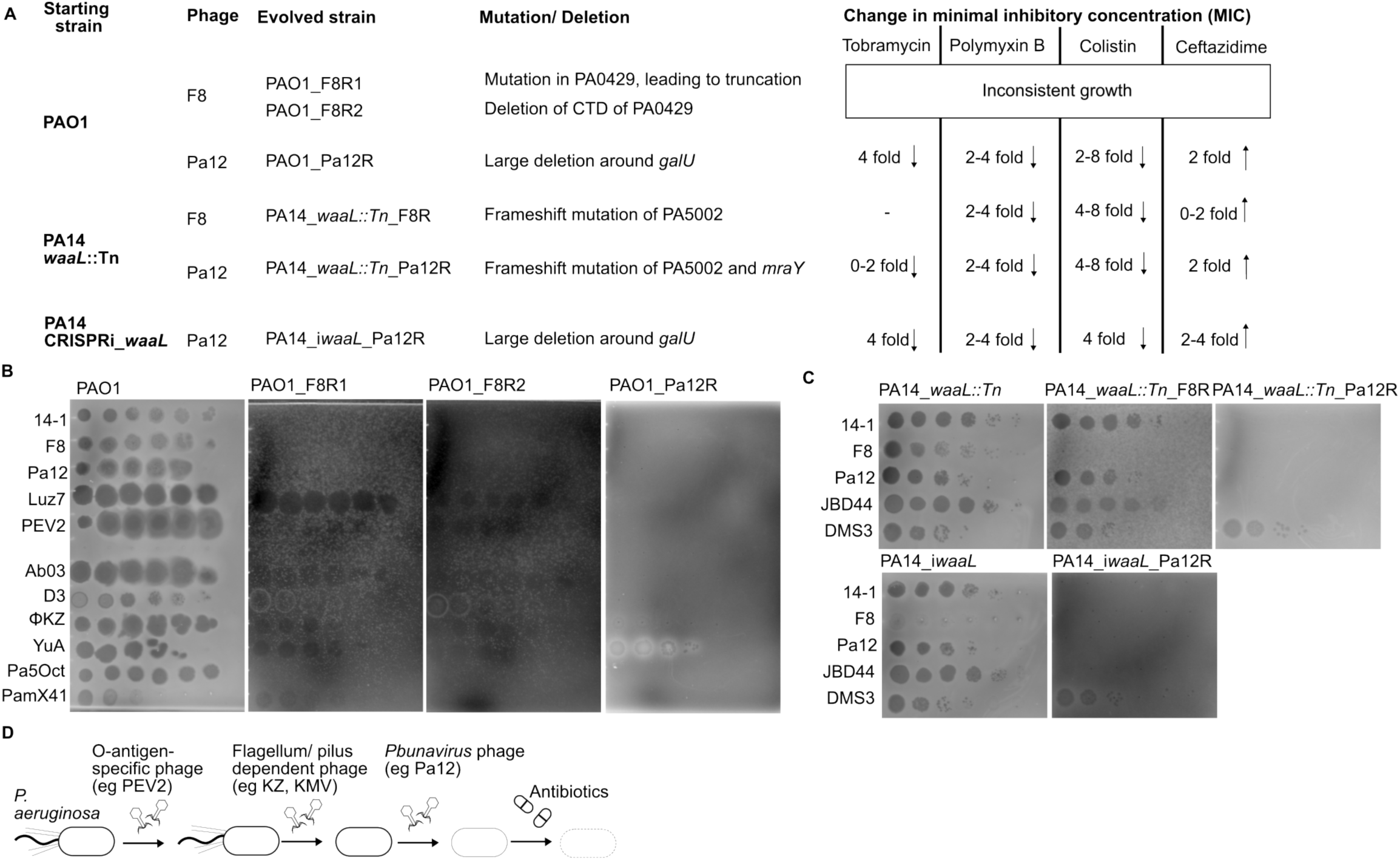
Evolved resistance against Pa12 phage confers resistance across many distinct phages. (A) Schematic of evolving resistance against F8 (“F8R”) or Pa12 (“Pa12R”). The characterized genetic mutation of each strain compared to parental PAO1 or PA14 and changes in MIC of each evolved phage resistant strain relative to parental PAO1 or PA14 (n=3) in cation-adjusted Mueller-Hinton Broth. (B) Plaque assay of PAO1 and evolved phage resistant strains with different LPS/pilus/flagellum-dependent phages. (C) Plaque assay of PA14 and evolved phage resistant strains with JBD44, DMS3, 14-1, F8 and Pa12. (D) Proposed sequential phage therapy treatment.

In the former, F8-resistant clones derived from PAO1 mutants (PAO1_F8R1 and PAO1_F8R2) carried mutations and deletions in PA0429 (Figure 6A). Analysis of PA0429 revealed sequence and structural homology with UDP-glucosyltransferases from other bacteria, suggesting a role in LPS biosynthesis (Figure S6). F8-resistant PA14 *waaL::Tn* acquired a frameshift mutation in *EIP97_27935*, which encodes for a PIG-L deacetylase family protein (PA5002 in PAO1; *dnpA*), while a Pa12-resistant derivative of the same background acquired frameshift mutations in both *dnpA* and *EIP97_RS09810,* annotated as a *mraY* family glycosyltransferase. We next tested the growth and antibiotic resistance of the evolved strains in cation-adjusted Mueller-Hinton broth (CAMHB), the suggested media for testing antibiotic resistance^36^. We observed that even in the absence of antibiotics, F8-resistant PAO1 mutants exhibited substantially reduced growth fitness and inconsistent growth kinetics. The F8/Pa12-resistant derivatives of PA14 *waaL::Tn*, however, grew well and showed a 2–8-fold decrease in minimum inhibitory concentration (MIC) for polymyxin B, and colistin, no/slight decrease in tobramycin MIC and a slight increase in ceftazidime MIC (Figure 6A). We further assessed whether *Pbunavirus* resistance conferred cross-resistance to other phages, a critical consideration for sequential phage therapy. F8-resistant mutants with PA0429 mutations retained susceptibility to non-*Pbunavirus* LPS-dependent phages, indicating that the underlying LPS modifications are specific enough to spare other receptor interactions (Figure 6B). F8-resistant derivatives of PA14 *waaL::Tn* only conferred resistance to F8, but PA14 *waaL::Tn_*Pa12R conferred resistance to JBD44, an LPS-dependent phage, but not DMS3 (Figure 6C).

Two Pa12-resistant mutants in both PAO1 and PA14_*iwaaL* backgrounds acquired large deletions encompassing the *galU* locus (Figure 6A), mirroring findings from independent studies^30,32^ and confirming the reproducibility of this evolutionary trajectory. Pa12-resistant mutants in both PAO1 and PA14 showed a 2–8-fold decrease in MIC for tobramycin, polymyxin B, and colistin. A slight increase in ceftazidime MIC was observed (Figure 6A). We observed that both Pa12-resistant mutants carrying *galU* deletions were resistant to all tested phages, including the jumbo phage ϕKZ, with the exception of the pilus-dependent YuA and DMS3 (Figure 6B-C). Therefore, results of Pbunavirus resistance across two different phages and across both PAO1 and PA14 reinforces the pattern that LPS alterations selected for by *Pbunavirus* resistance spares pilus-dependent phages.

These findings have direct implications for phage therapeutic sequencing: *Pbunavirus* phages should ideally be deployed after O-antigen-dependent phages or ϕKZ to avoid cross-resistance, and evolved *Pbunavirus* resistance predictably maintains or creates a vulnerability window for pilus-dependent phages or aminoglycoside antibiotics, respectively, both of which should be applied after effective *Pbunavirus* treatment to exploit this evolved sensitivity (Figure 6D).

## Discussion

A prevailing assumption has been that phage host range is governed by receptor presence or absence, and that resistant isolates often lack the required surface target. Our identification of L-Rha on the LPS inner core as the conserved *Pbunavirus* receptor encouragingly suggests that the molecule is present on most *Pseudomonas* isolates, albeit often concealed beneath by the O-antigen. Reliance on a conserved core component rather than the variable O-antigen likely underpins the characteristically broad host range of this family. Previous reports of O-antigen specificity^19,37^ can be reconciled by recognizing that different serotypes present barriers of varying stringency, which some tail fibers penetrate more readily than others. Critically, surface access to L-Rha is context-dependent: uncapped LPS upregulation in minimal medium masks the receptor even in strains that are fully susceptible in rich media, cautioning that standard *in vitro* susceptibility testing may overestimate therapeutic efficacy. Given that *migA* expression is elevated in lung environments^38^, this concern extends directly to the clinic, where *Pbunavirus* candidates for respiratory phage therapy should be screened for infectivity under conditions that reflect pulmonary uncapped LPS upregulation, not just standard laboratory media.

Identifying the receptor, however, solves only the first barrier. Despite intense interest in cataloguing antiphage defense systems through computational approaches, their direct contribution to phage therapy outcomes has remained unclear. Our results demonstrate that intracellular defenses play an important role in phage resistance, as single-step infection assays revealed that approximately 27% of clinical isolates permitted *Pbunavirus* adsorption but blocked replication, which is a frequency comparable to the 30% that resisted at the surface. A third group of strains emerged where wild-type phages replicate successfully, but deletion of the phage anti-defense island sensitizes the phage to intracellular defenses that are silently neutralized (by gp80, gp82, or gp88), suggesting that the true prevalence of active intracellular barriers is higher still. Previous research has shown that the anti-defense island encodes for inhibitors against multiple DNA-targeting defense systems including Zorya type I, RADAR, Hypnos, Druantia type I and the END nuclease^9,10^. While F63912 carries an intracellular barrier that can be inhibited by gp80 and gp82, sequence analysis of the strain shows absence of the aforementioned defense systems, suggesting that gp80 could have a larger range than previously described. We additionally reveal that gp82 is an important anti-defense gene in *Pbunavirus* phages against intracellular barriers in clinical isolates. Overall, these findings establish that majority of the clinical isolates in our panel harbor functional intracellular defenses against Pbunaviruses, and the phage anti-defense locus provides solutions for some but not all of these systems. Overcoming or bypassing the remaining barriers remain an open challenge for the future.

These findings converge on a specific phage therapeutic sequence. A phage that can access L-Rha, utilize it for infection, and replicate efficiently thanks to a sufficient complement of defense anti-defenses will impose strong selective pressure on the host. *Pbunavirus* replication pressure drives evolution toward core LPS truncation, particularly *galU* deletion, which eliminates L-Rha and confers cross-resistance to nearly all LPS-dependent phages simultaneously. However, this resistance mechanism carries substantial fitness costs: *galU* mutants exhibit increased sensitivity to antibiotics and display growth defects, while retaining full susceptibility to pilus-dependent phages. Additionally, loss of O-antigen is often associated with decreased virulence^39,40^. This stereotyped and exploitable resistance outcome argues for a defined therapeutic sequence: O-antigen-specific phages should be deployed first, as resistance through O-antigen loss exposes L-Rha and without affecting sensitivity to ϕKZ while sensitizing bacteria to Pbunaviruses; a ϕKZ-like phage should then follow, as resistance to ϕKZ-like phages usually does not drive LPS selection^41,42^. Pbunaviruses should then be used, driving core LPS loss; and finally, antibiotics can exploit the resulting vulnerabilities. Because effective *Pbunavirus* phage infection induces broad cross-resistance to other LPS-dependent phages, Pbunaviruses should be positioned last among this class. If pressure from the ϕKZ-like phage was incomplete and the type IV pilus is maintained, a pilus-dependent non-φKZ-like phage can be used as a last resort in tandem with, or after, a *Pbunavirus*.

In summary, we establish a layered framework for understanding and exploiting *Pbunavirus*-host interactions. This framework of identifying causal barriers at each level, engineering or recruiting solutions, and anticipate evolutionary consequences should guide the rational design of next-generation phage therapeutics.

## Methods

### Plaque assay

Bacteria strains were first grown overnight. 100-200 μl of overnight was then mixed with 0.4% agar before putting on plates with/without the appropriate inducers. The agar was left to dry, before 2 μl of serially diluted phages in SM buffer were spotted on. Once the spots dried, plates were incubated at 37°C. For screening assays, a strain was determined to be resistant to *Pbunavirus* if EOP < 10^-4^ compared to PAO1.

### Bacteria strains construction

pHERD30T (shortened to p30T) cloning was carried out as per previous. Briefly, the p30T/p20T backbone was amplified using primers WX_16/WX_17. Inserts were amplified using the respective primers (Table S1). Gibson assembly was then done and incubated at 50°C for at least 1h. Gibson mixture was then transformed into XL1blue using heat shock and selected on either gentamicin (p30T) or carbenicillin (p20T). Colonies were then selected and sent for sequencing with Quintara Biosciences. Plasmids were then introduced into PAO1 using electroporation. Separately, plasmids were introduced into PA14 and other clinical isolates by first electroporating into *E. coli* SM10 cells, then conjugated into *P. aeruginosa* strains by mixing *E. coli* and *P. aeruginosa* and left to grow for at least 4h. Transformed *P. aeruginosa* was then selected using VBMM in the presence of either gentamicin (p30T) or carbenicillin (p20T).

To make deletion strains in PAO1 and PA14, homologous recombination constructs were first cloned into the PMQ30 plasmid. Constructs were then either electroporated into PAO1 or SM10. For PA14 deletion, SM10 carrying PMQ30 plasmid and PA14 was mated to introduce the plasmid in. *P. aeruginosa* with PMQ30 plasmids integrated were then selected on plates with gentamicin, and counter-selected in the presence of sucrose. Individual colonies were then picked and checked for gentamicin resistance before PCR or sequencing was carried out to confirm the deletion. Strains were then stored in glycerol at -80°C.

### Phage engineering

To delete or change genetic loci in phages, homologous recombination was used. Templates for recombination were first cloned into the p30T plasmid after linearization with NheI enzyme, using Gibson assembly. Gibson mixture was then transformed into XL1 blue, before selecting on gentamicin and sequencing with Quintara Bio. Verified plasmids were then transformed into PAO1 as described above. Approx. 100-1000 PFU of the target phage (eg. F8, 14-1, Pa12) were then used to infect PAO1 carrying the plasmid for recombination. After overnight growth, phages were harvested using chloroform treatment. Recombinant phages were then selected and grown in the presence of CRISPR-Cas IF (Cascade Cas3) selection.

### Growth curves

PAO1 carrying CRISPRi guides were grown overnight in the presence of induction. Bacteria were then subcultured for 2h either in M9 + 1mM IPTG or LB + Mg^2+^ + 1mM IPTG prior to addition of phages at the MOI shown. Bacteria growth was then followed over time in a 96-well plate using a H1 synergy plate reader.

### Single step phage assays

For single step assay to determine adsorption in PA14 background, strains were first grown in the presence of gentamicin in tubes. The next day, a 1:30 dilution was made in fresh media in the presence of arabinose and grown for approx.. 2-3h until log phase growth was observed on the plate reader. Phages were then added and samples were taken at t = 15 and added directly to chloroform. The aqueous layer was then collected for plaque assays.

To determine if clinical isolates carried intracellular or surface defenses, all strains were first grown in a deep well 96 well plate. The next day, a 1:20 dilution was made in fresh media and grown for approx. 2-3 h until log phase growth was observed on the plate reader. Phages were then added, and samples were taken at t = 15 or t = 90 min and prepared with chloroform as described above.

Finally, to determine if internal defenses are abortive systems, t = 120 instead of t = 90 was used instead to allow for maximum effect on bacteria to be seen (in case there is a delay in the effect on the bacterium). MOI > 1 was also used to check for abortive phenotype. In addition to phage collection described above, bacteria pellet was also collected after centrifugation at 10,000 x g. The bacteria pellet was then washed 1x with PBS, before resuspending in LB, serially diluted and 5 μl plated on LB plates. Bacteria CFU and phage PFU per well was then calculated.

### Minimal inhibitory concentration (MIC) assay

Bacteria preparation: To adhere to clinical standards, MIC testing was carried out using Cation-adjusted Mueller-Hinton Broth (CAMHB). Briefly, bacteria were streaked out overnight on LB + Mg^2+^ plates and grown for <18 h. Bacteria were resuspended in CAMHB and OD_600_ was measured, before diluting to approx. 10^5^ CFU per well (2x bacteria concentration)^36^.

Plate preparation: Antibiotic stocks were prepared fresh the day of usage. Antibiotics were then added to CAMHB to a final concentration of 2x the antibiotic test concentrations, and then serially diluted 2-fold. Bacteria suspension and the 2x antibiotic CAMHB were then mixed. Bacteria were then grown overnight in BioTek LogPhase 600 plate reader (Agilent) reader for 24h and an OD600 reading was taken every 10 minutes. A no antibiotic control was included to check for baseline growth in the absence of antibiotics. Antibiotic concentrations tested ranged from 0.125 – 8μg/ml.

### Whole genome sequencing

Genomes were extracted using the modified SDS/Proteinase K method. Briefly, bacteria pellet was resuspended in 100ul of water and mixed with lysis buffer. For phages, 100 μl of phage lysate was used instead. The final concentration of the lysis buffer was 10 mM Tris, 1 mM EDTA, 0.5% SDS. The solution was then incubated at 37 °C for 30 min, followed by 55 °C for 30 min-1 h. DNA was then purified using Genomic DNA Extraction kit and quantified with Nanodrop. Sequencing of bacterial isolates was carried out using Plasmidsaurus. Sequencing of phage genomes was carried out in-house on a MiSeq and MiSeq i100, at the UCSF CAT. Breseq and Snapgene alignments were used to identify mutations.

### qPCR

After whole genome extraction as described above, approx. 1-5ng of DNA was then used for qPCR with Luna Universal qPCR Mastermix (NEB), with primers WX_Q21F and WX_Q21R for bacteria (described for *Pseudomonas aeruginosa* previously^43^) and primers WX_Q2F and WX_Q2R for Pa12, on the CFX Connect Real-Time PCR Detection System (Bio-Rad).

### CRISPRi library construction

#### IPTG-inducible *P. aeruginosa* CRISPRi system construction and characterization

An IPTG-inducible, mobilizable, chromosomally-located CRISPRi system for *P. aeruginosa* was developed by modifying previously constructed Mobile-CRISPRi systems with swappable modular components and gene-targeting spacer sequences cloned into a BsaI site (see Figure S7A and Tables S1-3)^44–46^. The sgRNA was expressed from a variant of the *trc* promoter without a lac operator and a codon-optimized dCas9 (*Hsa* Spy::3Xmyc dCas9) was expressed using an IPTG-inducible synthetic promoter ’vD’^47^. CRISPRi knockdown strains were constructed by transferring a Tn*7* transposon harboring the CRISPRi system components by conjugation and transposition from *E. coli* pir^+^ DAP^-^ donors to the *P. aeruginosa* recipient with selection for chromosomal integration with gentamicin (30 µg/ml) LB-agar plates and counterselection against the *E. coli* donors by omission of diaminopimelic acid (DAP). A strain with the CRISPRi system was assayed to determine that they had no growth defect compared to WT by monitoring growth at OD_600_ with and without induction at 1mM IPTG in a Tecan Mplex plate reader for 20h (see Figure S7B). CRISPRi knockdown of ∼10-fold was quantified by assaying cells passaged ∼15 generations with 1mM IPTG induction using a fluorescent reporter (sfGFP) expressed from the transposon and a guide targeting this gene (see Figure S7C). Cells were washed with 1 x PBS before fluorescence was measured using a Tecan Mplex plate reader and normalized to cell density.

#### *P. aeruginosa* genome-scale CRISPRi library construction

A library of 30,934 guide sequences consisting of ∼4 perfect match guides/gene (full knockdown), ∼10 mismatch guides/essential gene (gradient of partial knockdown), and ∼1000 non-targeting controls was designed using custom Python scripts^46,48^. Guides (Table S2) were selected to target both *P. aeruginosa* PA14 and/or PA01 genomes, and essential genes were selected based on the *Pseudomonas* core essential genome defined by Poulsen, *et al*.^49^. The genome-scale library was constructed by amplification of the sgRNA spacer-encoding sequences from an oligo pool (Agilent Sure-Print custom pool OL-I-2020; see table S3) with primers oJMP1184 and oJMP1185 and Q5 DNA polymerase (NEB), ligation of the amplicon digested with BsaI-HF-v2 (NEB) into the BsaI site of the transposon-harboring plasmid (pJMP6777), and transformation into the *E. coli* mating strain with selection of ∼2-3 million CFUs on LB + carbenicillin (100µg/ml) + DAP (300µM) agar plates which were harvested and stored in LB + 15% glycerol at -80°C. Tri-parental conjugation of this library donor strain (sJMP6828) along with the Tn7-transposase donor strain (sJMP2954) and either the *P. aeruginosa* PAO1 or PA14 recipient strain (sJMP742 or sJMP744, respectively) was carried out for 5 hours at 37°C on an LB + DAP (300 µM) agar plate, followed by selection of ∼1-2 million transconjugants on LB + gentamicin (30 µg/ml) agar plates. The CRISPRi system contained on a Tn7 transposon is thereby stably inserted on the chromosome in the *att*_Tn7_ site downstream of *glmS*^50^. Finally, the libraries were harvested and stored in LB + 15% glycerol at -80°C (PA14: sJMP10551 and PAO1: sJMP10587).

### CRISPRi analysis

For PA14 CRISPRi analysis, library was defrosted from -80°C into 50 ml at OD_600_ = 0.03. Library was then grown in the absence of induction till OD_600_ = 0.2. Library was then subcultured in 1:10 and grown in the presence of induction overnight. Library was then diluted to OD_600_ = 0.2 in the presence of inducer and grown until OD_600_ reaches 0.4. Phages were added at an MOI of approx. 50 for 14-1. No phage control is present. Infection was tracked over time, and once the start of a crash is observed, approx. 600 μl bacteria were collected. Cultures were then split into 2, washed 1x with PBS and then immediately frozen on dry ice. Pellets were kept in -20°C. For PAO1 CRISPRi analysis, library was induced overnight and then diluted to OD_600_ = 0.1 in minimal media and inducer the next day and grown for at least two hours until OD_600_ = 0.2. F8 was then added at an MOI of approx. 5.

Genomic DNA extraction was carried out using Qiagen kit according to manufacturer’s instructions, as per previous^51^. CRISPRi guides were amplified using primers 697/698. i7 and i5 indexes were then added using the Ultra II Q5 polymerase (NEB), using primers from NEB E6440S. Sequencing was done on the NovaSeq at the UCSF CAT, with 20-30% phiX. Analysis was done using heuristicount.py, followed by a script on R to plot changes in frequency of genes.

### Bioinformatics and statistical analysis

Analysis of defense systems was carried out using DefenseFinder using protein fasta sequences^52^. Guides for PAO1 and PA14 are featured on ryandward/Pseudomonas_sgRNA. Analysis of CRISPRi screens was carried out using heuristicount R script available on GitHub (ryandward/phylogenetic_CRISPRi). Output of frequency of each gene’s representation within the library were then input into R to plot changes in frequency. HHpred^53^ and AlphaFold3^54^ was used to analyze PA0429. For generation of efficiency of plating (EOP) bar graphs, GraphPad PRISM was used. For statistical analysis of number of defense system, the normality test followed by Mann-Whitney test on GraphPad PRISM was used. Biorender was used for drawing some schematics.

### Cryo electron tomography

*P. aeruginosa* cells (OD_600_ = 0.5) were mixed with purified phage 14-1 (5 mg/mL) at 1:1 ratio to a total volume of 12 µl and incubated at 37 °C for 10 minutes for phage adsorption. One µl of 10 nm Protein-A-gold fiducials (CMC Utrecht) was added to the 12 µl sample. Three µl of the sample was then applied to a freshly glow discharged Quantifoil R3.5/1 Cu/Rh 200 mesh grid, incubated for 30 s, and then manually blotted via blotting with a filter paper (Electron Microscopy Sciences) from the back of the grid and fixed with 2.5 μL 2% paraformaldehyde for 1 minute. The sample was reblotted for 5 seconds (total force -10, wait time 10 seconds) and plunge-frozen into liquid ethane in the Vitrobot IV (Thermo Fisher Scientific), while the blotting chamber was maintained at 100% humidity at 10 °C. The grids were clipped and stored under liquid nitrogen until cryo-ET data collection was performed.

The data was collected on a Titan Krios G3 microscope (Thermo Fisher) operating at an acceleration voltage of 300 kV, fitted with a Quantum energy filter (slit width 20 eV) and a K3 direct electron detector (Gatan) using the SerialEM software^55^. Cryo-ET tilt series were collected using a grouped dose-symmetric tilt scheme as implemented in SerialEM^56^, with a total dose of 110 electrons/Å^2^ per tilt series, defocus range of between −3 to -5.5 μm target defocus, and with ±60° tilts of the specimen stage at 3° tilt increments. Tilt series images were collected using a physical pixel size of 2.67 Å.

Cryo-ET data processing was performed using RELION5.0^57^. After tilt series import, tilt-series movies were motion-corrected and split into odd and even images with MotionCor2^56^ implemented in RELION-5.0. Tilt-series alignment and initial contrast transfer functions (CTFs) estimation were performed using ARETOMO2^58^. Tomograms were generated using RELION5.0 and denoized with Cryo-CARE^59^. Figure panels containing cryo-ET images were prepared using IMOD^60^ and Fiji^61^.

## Acknowledgements

J.B.-D. is supported by the National Institutes of Health (nos. R01 AI171041 and R01 AI167412). This work is also partly supported by the NUS Development Grant awarded to W. X. Y. T. A. M. B. is supported by the Medical Research Council, as part of United Kingdom Research and Innovation (also known as UK Research and Innovation) [Programme MC_UP_1201/31] and the Wellcome trust (grant 225317/Z/22/Z). We would also like to thank William J. Heelan for technical assistance in this project.

We would also like to thank Brouns lab at TU Delft for the vB_PaeM_FBPa10 and vB_PaeM_FBPa12 phage (annotated in this paper as Pa10 and Pa12 respectively). Sequencing was performed at the UCSF CAT, supported by UCSF PBBR, RRP IMIA and NIH 1S10OD028511-01 grants.

## Author contributions

X. Y. - Investigation, Conceptualization, Methodology, Validation, Visualization, Writing - original draft, Writing - review and editing

B. B. - Investigation, Methodology, Writing - review and editing

D. W. - Investigation

M. - Investigation

L. - Investigation

H. - Investigation

G. - Investigation, Visualization

C. L. - Investigation

A. M. B. - Funding acquisition, Investigation, Writing - review and editing

M. P. - Writing - review and editing

B. D. - Conceptualization, Funding acquisition, Investigation, Writing - original draft, Writing - review and editing

## Supplementary Figure legends

**Figure S1:**
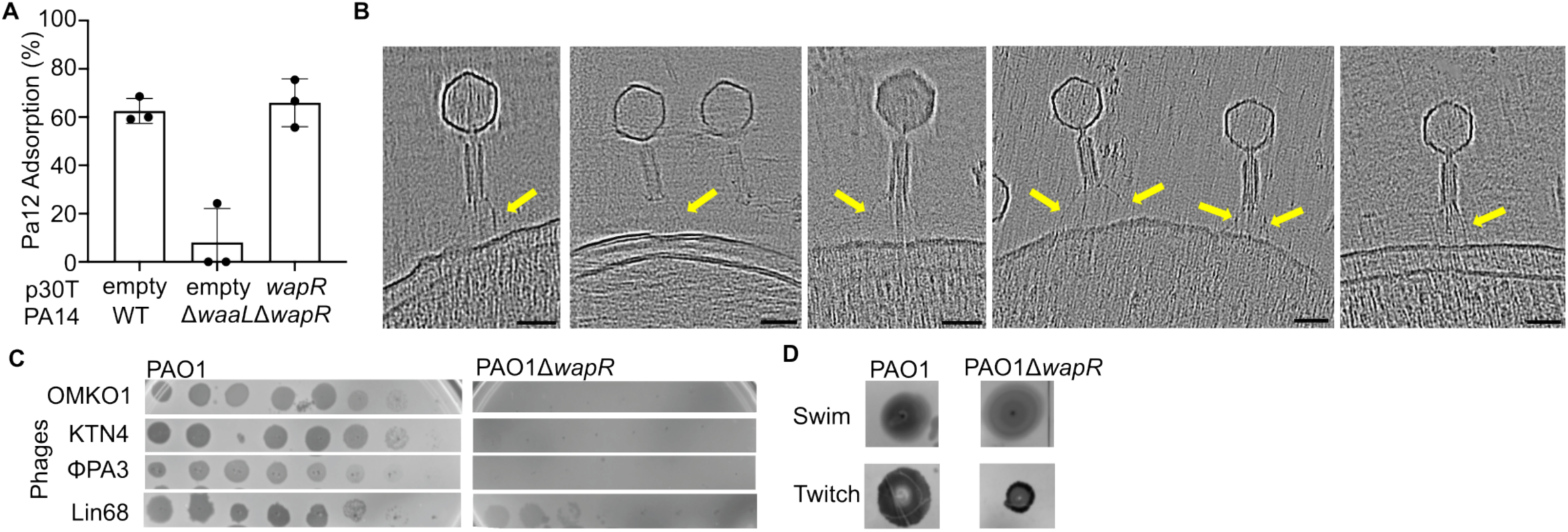
Phages use the core LPS as receptor. (A) Adsorption assay of Pa12 on PA14 + p30T empty, PA14Δ*waaL*Δ*wapR* + p30T empty, PA14Δ*waaL*Δ*wapR* + p30T *wapR*. (B) A gallery of slices from cryo ET data of 14-1 phage infecting PAO1 cells. Yellow: phage tail fibers extending close to the outer membrane. Red: phage sheath orthogonal to the outer membrane, presumably in the act of injection. Scale bar: 50nm. (C) Plaque assays of related jumbo phages on PAO1 with/without *wapR*. (D) Swimming and twitching assays for PAO1 and PAO1Δ*wapR*.

**Figure S2:**
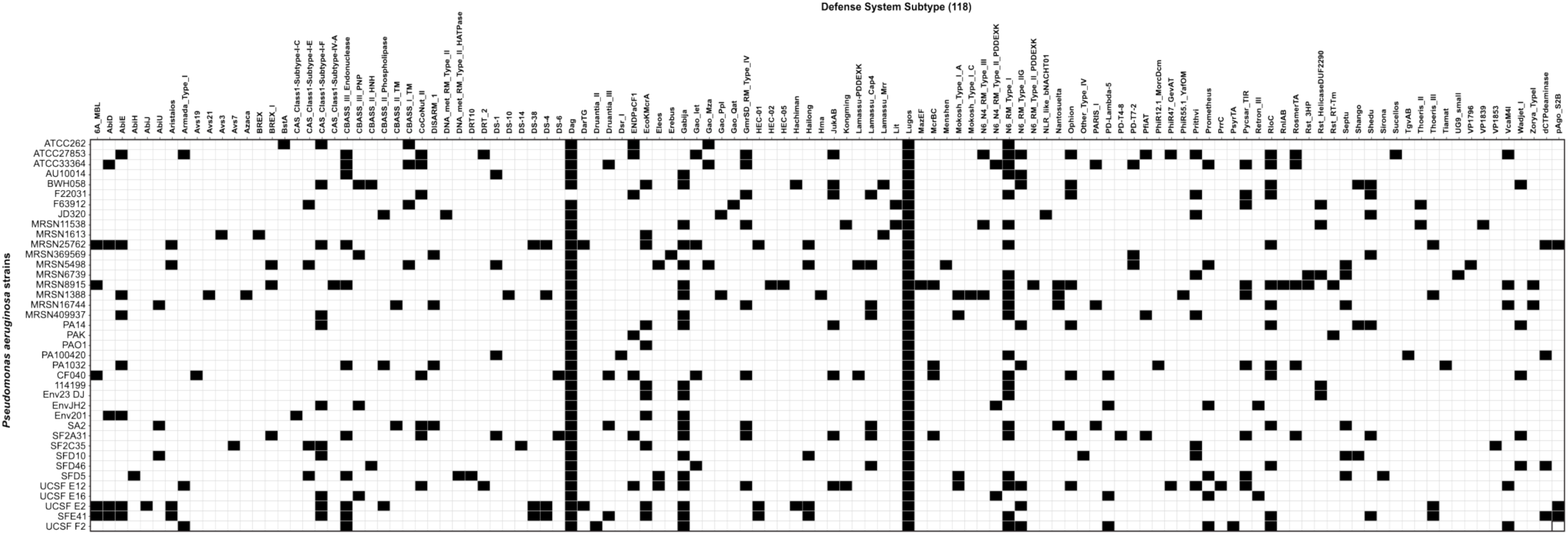
Presence of different characterized defence systems in clinical isolates tested in this paper. Defense systems were detected using HMM-based sequence homology search via DefenseFinder

**Figure S3.**
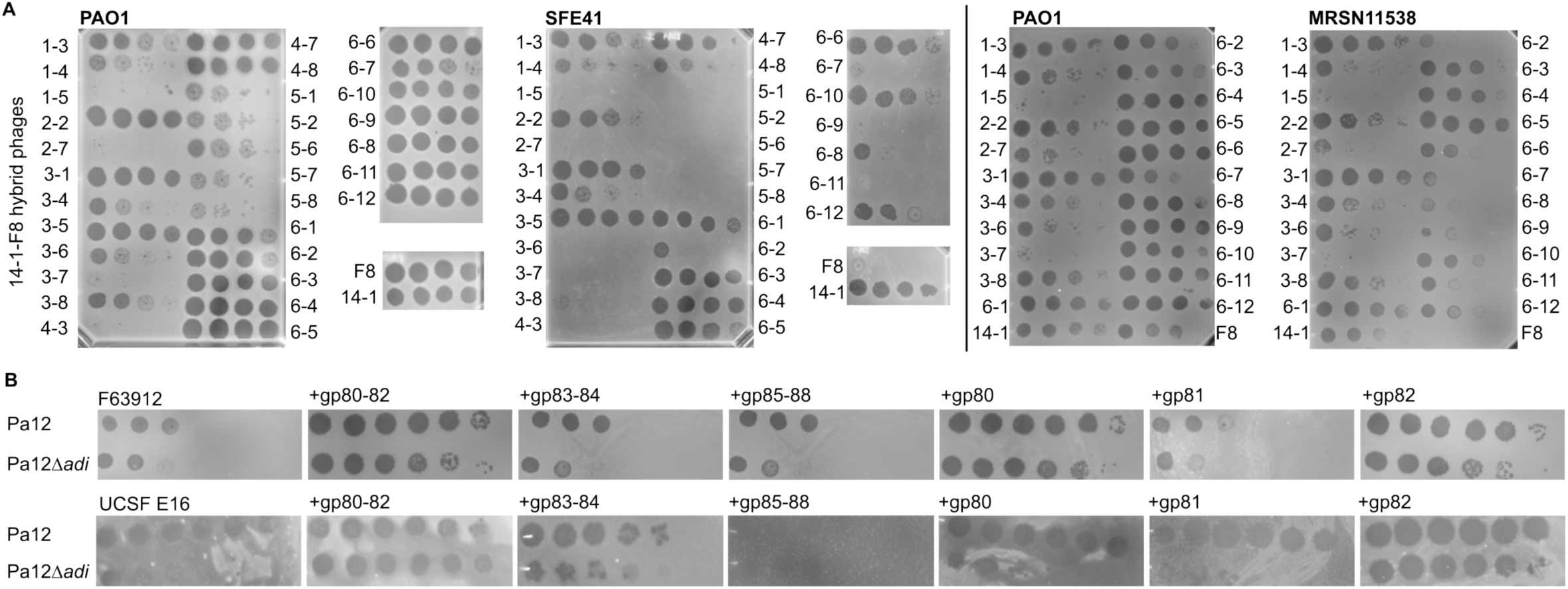
Identifying phage solutions to barriers in clinical isolates. (A) Plaque assay of 14-1, F8 and 14-1-F8 hybrid phages (shown as numbers) on SFE41 and MRSN11538. (B) Plaque assay of Pa12 and Pa12Δ*adi* on F63912 and UCSF E16, with different genes from phage Pa12 cloned.

**Figure S4.**
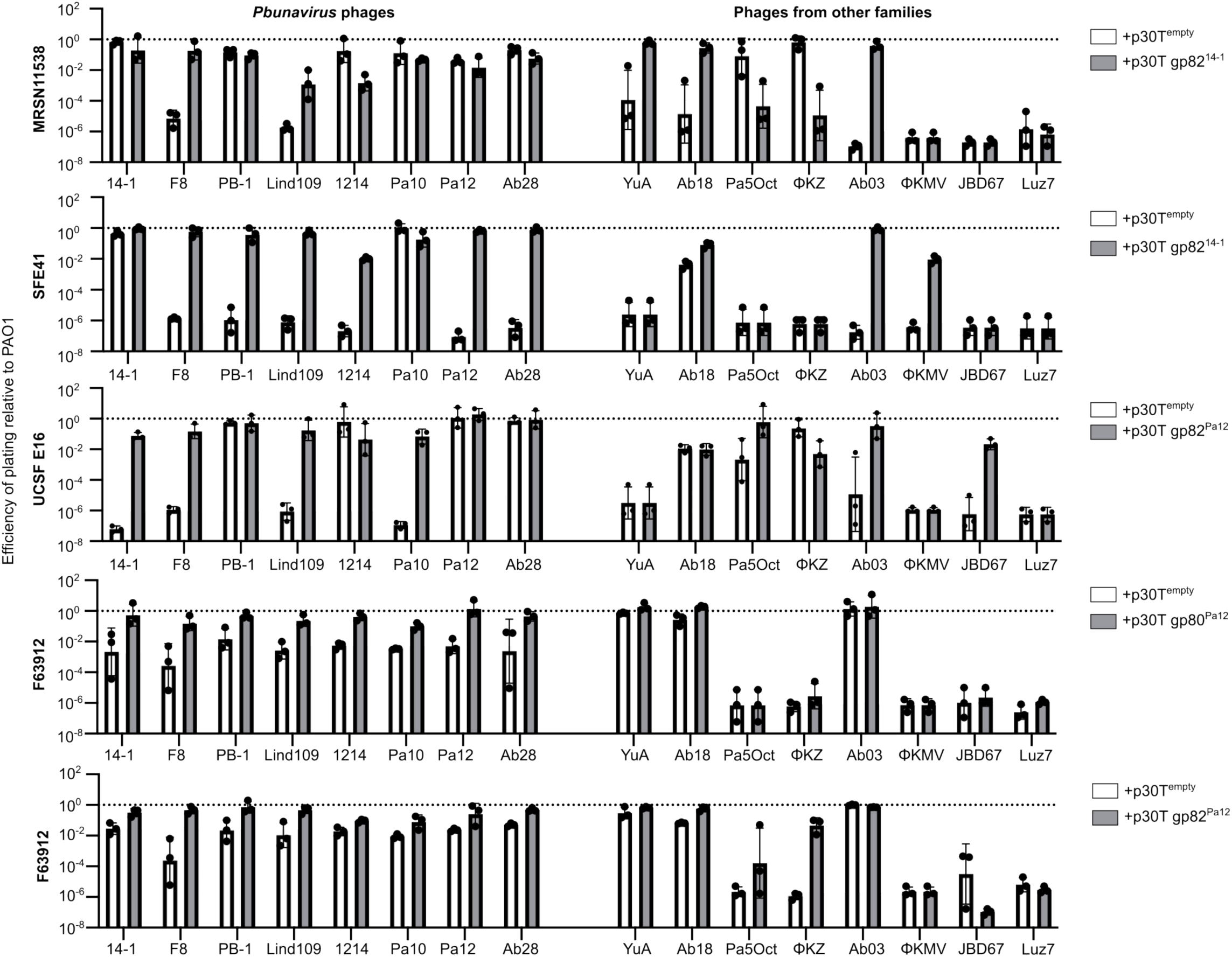
Efficiency of plating (EOP) of phages from different families on F63912, UCSF E16, SFE41 and MRSN11538 compared to PAO1 in the absence/presence of anti-defenses identified in Pa12/14-1.

**Figure S5.**
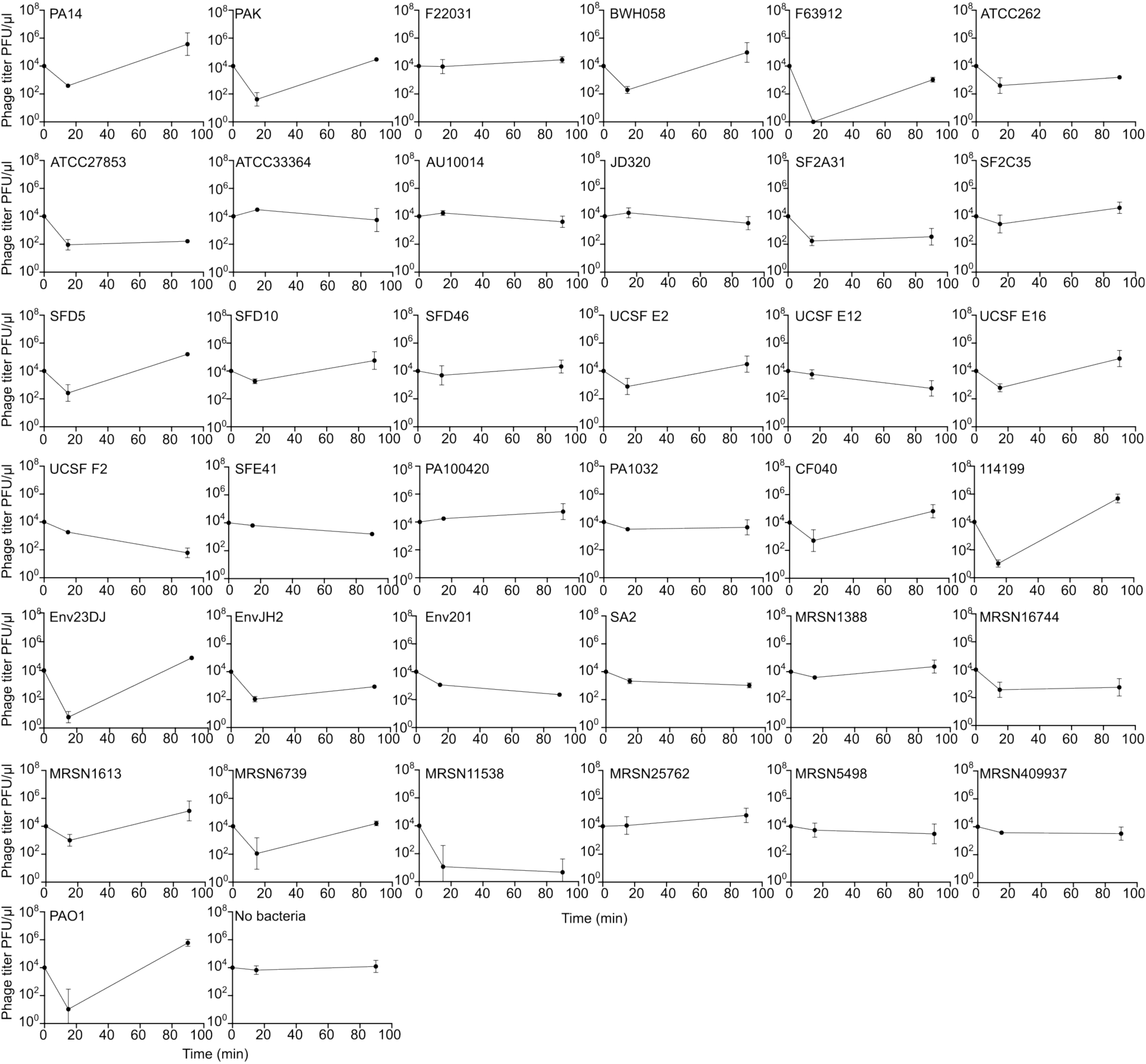
Single step growth curve of Pa12 phage infection across different clinical isolates.

**Figure S6:**
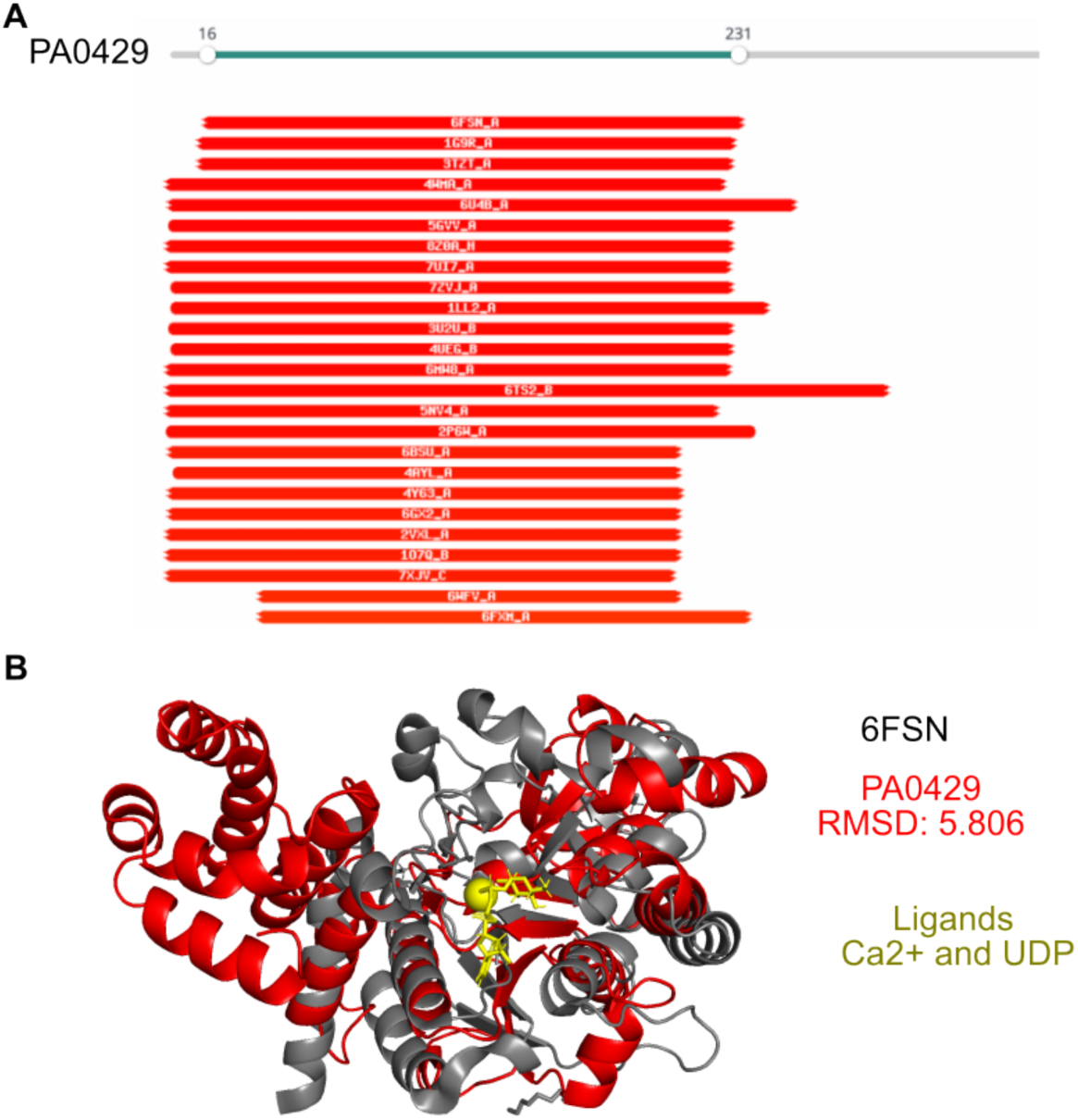
PA0429 shares homology with UDP glucosyltransferases. (A) HHpred analysis of PA0429. (B) AlphaFold prediction of PA0429, and overlay with characterized UDP glucosyltransferase 6FSN.

**Figure S7:**
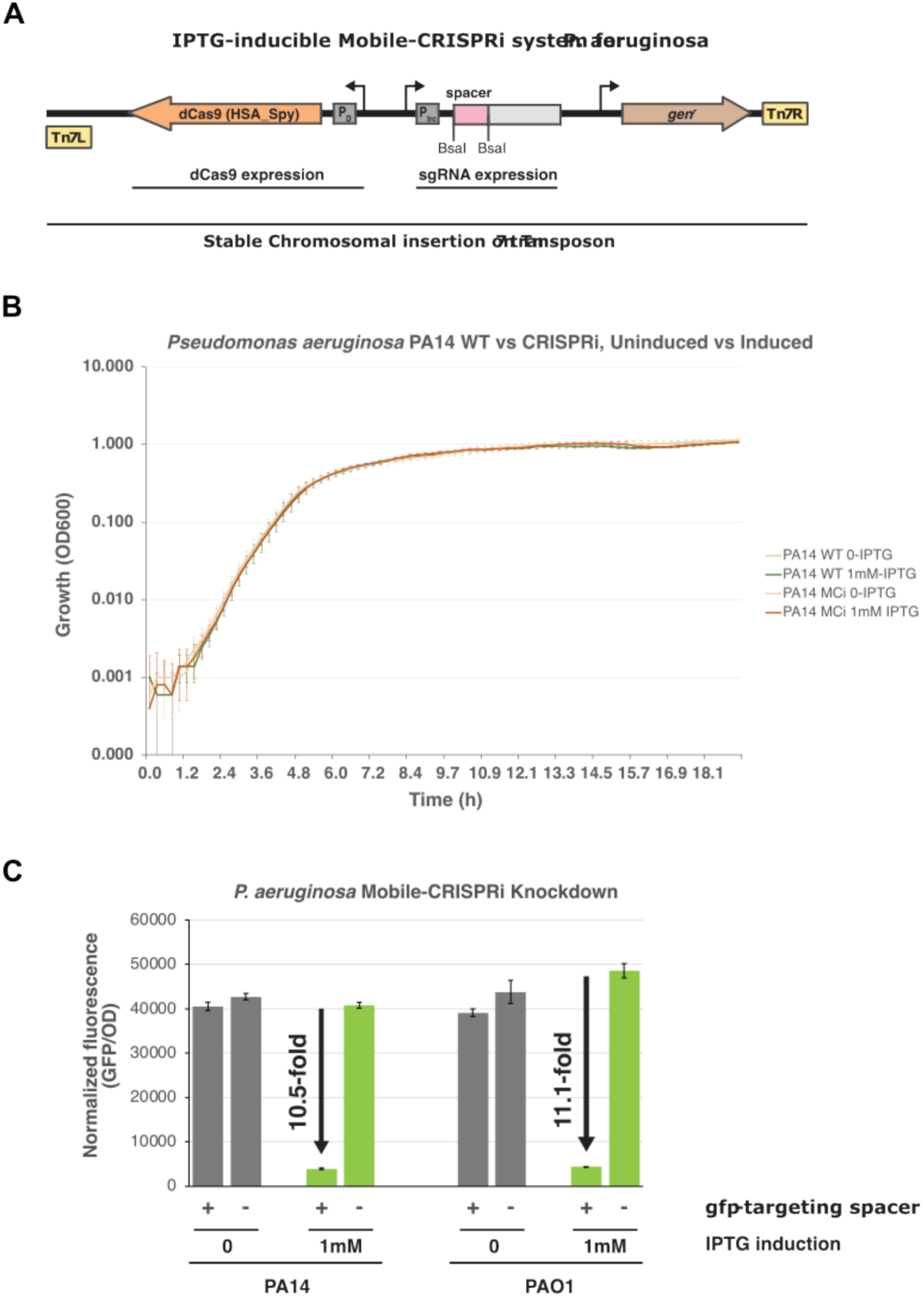
Mobile-CRISPRi validation in *P. aeruginosa*. (A) Schematic for IPTG-inducible mobile-CRISPRi system for *P. aeruginosa*. (B) Growth curve of PA14 WT or a strain carrying a mobile-CRISPRi system (MCi) in the absence/ presence of IPTG inducer. (C) Fluorescence quantification of CRISPRi knockdown of a fluorescent reporter with/without sgRNA spacer.

